## Supplementary materials for "Individuals with ventromedial frontal damage have more unstable but still fundamentally transitive preferences"

**This PDF file includes:**

Supplementary methods
Supplementary results
Fig. S1
Supplementary references

### Supplementary methods

**Stimulus norming.** The artwork stimuli were paintings that were rated highly by participants in Vaidya and Fellows (1). The set B stimuli consisted of 5 paintings by Monet, which were all within the top 20 most highly rated paintings by those subjects. We selected Monet as he was the artist that occurred most frequently in the top 20 rated paintings of Vaidya and Fellows (1). The 5 selected paintings were roughly similarly preferred (i.e., chosen with close to the same frequency in pair-wise choices across the whole sample) in a sample of 107 participants recruited from Amazon Mechanical Turk. Set A consisted of paintings of the similar style/era (Impressionist, Romantic periods) in the top 40 ranked paintings of the Vaidya and Fellows (1) stimuli set.

The chocolate bars were from five brands (Lindt, Godiva, Ghirardelli, Dove, and Cadbury). We selected five brands that were roughly similarly preferred across the population. These brands were being sold for similar prices, were rated similarly on a seven-point scale by a sample of 103 participants from Amazon Mechanical Turk (mean rating = 5.76), and were selected at roughly similar frequencies in pair-wise choices across another sample of 101 Mechanical Turk participants. Milk chocolate bars from each of the 5 brands were in set B, while dark chocolate and dark chocolate almond bars from each brand were in set A. The stimuli consisted of publicly available pictures of the front side of the chocolate bar packaging.

We used sixteen gambles of equal expected value ($8.80). The stimuli consisted of a pie chart showing the probability of winning, with text on top indicating both the cash amount to be won and the probability of winning. The five set B gambles were the “Cash II” set in Regenwetter, Dana and Davis-Stober (2), which used contemporary monetary equivalents of the Tversky (3) five gamble set. The probabilities were 28%, 32%, 36%, 40%, and 44%. Set A consisted of 11 other gambles with the same expected value (probabilities of 8%, 17%, 25%, 33%, 42%, 50%, 58%, 67%, 75%, 83%, 92%).

**Sensitivity of probabilistic tests.** We performed several simulations to determine the sensitivity of tests of the mixture model, i.e., the rate at which this test would declare different forms of random or heuristic-based choice to be intransitive. First, following Regenwetter, Dana and Davis-Stober (2), we randomly picked a choice probability for every pair from a uniform distribution (from 0 to 100%). As previously shown in that paper, only about 5% of the choice datasets simulated in this manner satisfy the triangle inequalities. That is, only 5% of the possible set of choice proportions for 10 pairs/5 stimuli satisfy the mixture model.

Second, we simulated an intransitive chooser who has an entirely consistent preference within each pair (i.e., choosing A 100% of time when it is paired with B) that is unconstrained by any higher order transitive structure (i.e., the preference in each pair is independent from that of all other pairs). This type of intransitive chooser only satisfies the triangle inequalities about 12% of the time for choice proportions for 10 pairs/5 stimuli.

Third, we simulated an intransitive chooser using the lexicographic semiorder heuristic (3). This heuristic is easiest to demonstrate with the gambles stimulus set. Following Tversky (3), we defined our lexicographic semiorder rule as follows: if two gambles are adjacent (i.e., next to each other in the set in terms of probabilities/payouts), always choose the gamble with the higher payout (amount); for all other (non-adjacent) gamble pairs, always select the gamble with the higher probability. Such a chooser would never satisfy the triangle inequalities in our dataset. Together, the first three sets of simulations show that our tests of mixture model are very sensitive to different forms of intransitive choice.

**Supplementary Results**

**Additional effects on DDM parameters.** As described in the main text, we performed a mixed ANOVA to test the effects of category (art, brands, gambles) and group on the DDM parameters. For the noise term, we found main effects of group ($F(2, 37) = 6.25$, $p = 0.005$) and category ($F(2,4) = 12.85$, $p < 0.001$), but no category x group interaction ($F(4,74) = 1.09$, $p = 0.37$). The main effect of group, as already explored in the main text, was driven by a higher noise term in the VMF group. The main effect of category was driven by a higher noise term in the art category (mean = 0.11) compared to both the brand (mean = 0.09) [$t(39) = 3.54$, $p = 0.001$] and gamble categories (mean = 0.08) [$t(39) = 3.96$, $p = 0.0003$)]. This result suggests that preferences in the art category were noisier across all participants.

For non-decision time, we found a significant main effect of category ($F(2,74) = 11.22$, $p < 0.0001$), but no significant effect of group, nor a significant category x group interaction ($F4,74) = 0.938$, $p < 0.45$). The main effect of category was driven by a longer non-decision time in the gamble category (gamble mean = 0.60) compared to both the art category (art mean = 0.35, $t(39) = -3.59$, $p < 0.0001$) and the brand category (brand mean = 0.36, $t(39) = 3.49$, $p = 0.001$). This result may be due to the fact that the gamble stimuli required more reading than the other two categories.

There were no significant effects of category, nor significant category x group interactions, for any other parameters in the DDM.

**Possible correlates of choice cycles.** We found that a subset of individuals with VMF damage show the most pronounced increase in choice cycles (overlap of their lesions are depicted in

**Figure S1**). However, we did not find evidence to support any particular account of this heterogeneity. The total number of choice cycles (i.e., across all three categories) was not significantly correlated with lesion size (in cc's), whether considering all subjects with lesions (Spearman's rho = -0.14, $p = 0.51$) or only those with VMF damage (rho = -0.13, $p = 0.67$). Within the VMF group, the total number of choice cycles was also not significantly correlated with lesion volume within a vmPFC mask defined based on value effects in fMRI studies (2) (rho = -0.06, $p = 0.83$). Finally, across all subjects, the total number of choice cycles was not significantly correlated with any demographic variables (gender, point biserial $r = 0.13$, $p = 0.39$; age, rho = 0.14, $p = 0.35$; education, rho = 0.24, $p = 0.11$

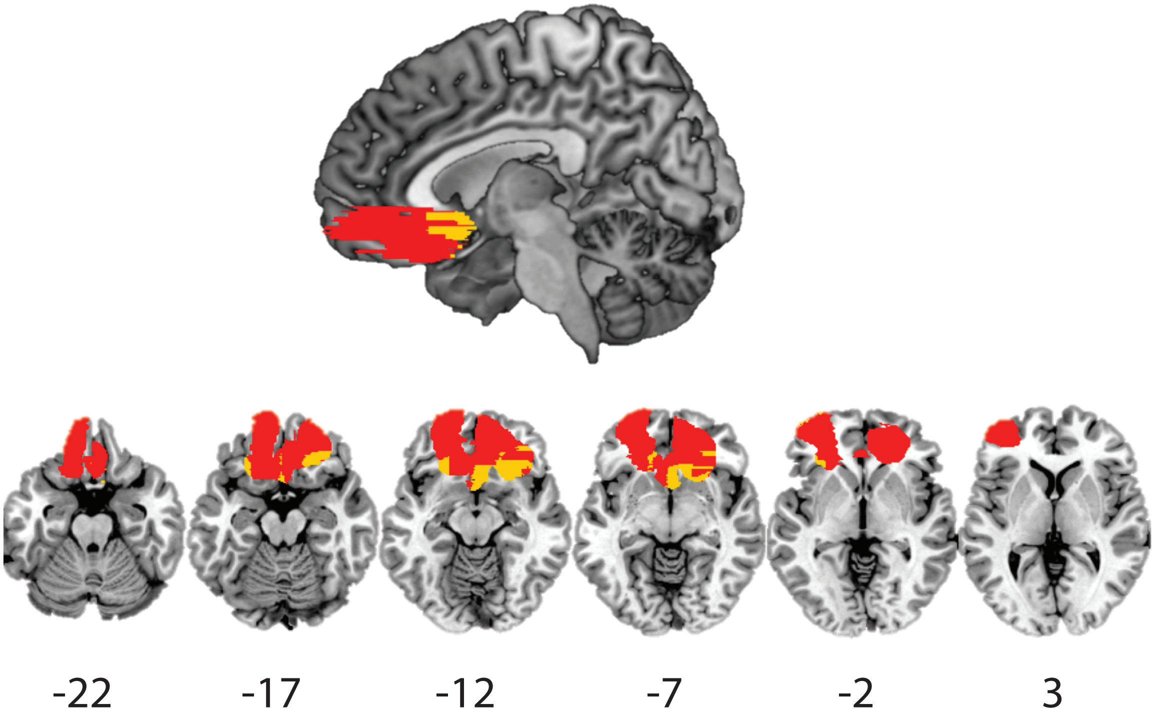


**Figure S1:** Lesion tracings of the three individuals with VMF lesions who had significantly more cyclical choices compared to healthy control subjects, as determined by case-control t-tests. Red denotes areas where at least one of these subjects had a lesion; yellow denotes the areas where at least one of these subjects had lesions *outside* of all other lesion subjects. There was very little overlap in lesions within the three subjects (only maximally two out of three and only in a small number of voxels). Numbers below axial slices indicate the MNI z-coordinates.

### Supplementary references

1. A. R. Vaidya, L. K. Fellows, Testing necessary regional frontal contributions to value assessment and fixation-based updating. *Nature Communications* **6**, 10120 (2015).
2. M. Regenwetter, J. Dana, C. P. Davis-Stober, Transitivity of preferences. *Psychological review* **118**, 42-56 (2011).
3. A. Tversky, Intransitivity of preferences. *Psychological review* (1969).
